# High prevalence and dependence of centrosome clustering in mesenchymal tumors and leukemia

**DOI:** 10.1101/2023.03.13.532472

**Authors:** N. Moreno-Marin, G. Marteil, N. C. Fresmann, B.P. de Almeida, K. Dores, R. Fragoso, J. Cardoso, J.B. Pereira-Leal, J.T. Barata, S. Godinho, N.L. Barbosa-Morais, M. Bettencourt-Dias

**Author notes:** Shared authors (G. Marteil, N. C. Fresmann, B.P. de Almeida).

## Abstract

The presence of supernumerary centrosomes is a hallmark of cancer and is frequently observed in aggressive tumors. Cancer cells with centrosome amplification achieve pseudo-bipolar spindles through specific coping mechanisms in order to survive. However, their distribution and prevalence in cancer remain largely unknown. Here, using the NCI60 panel of cancer cell lines, we show that the presence of coping strategies correlates with centrosome amplification, with the clustering of extra-centrosomes within the two spindle poles being the most widespread mechanism. Moreover, we report an association between centrosome clustering ability and the epithelial-to-mesenchymal transition (EMT) and observe that the induction of mesenchymal characteristics in breast cancer cells with centrosome amplification promotes clustering.

Furthermore, we unveil hematological malignancies, which lack epithelial characteristics, as the most proficient in centrosome clustering. Finally, we show that acute lymphoblastic leukemia is particularly sensitive to targeting clustering through inhibition of the spindle assembly checkpoint. Our study reveals how centrosome clustering and the EMT collaborate to promote carcinogenesis, suggesting new possibilities to treat tumors with low epithelial characteristics, in particular leukemias.

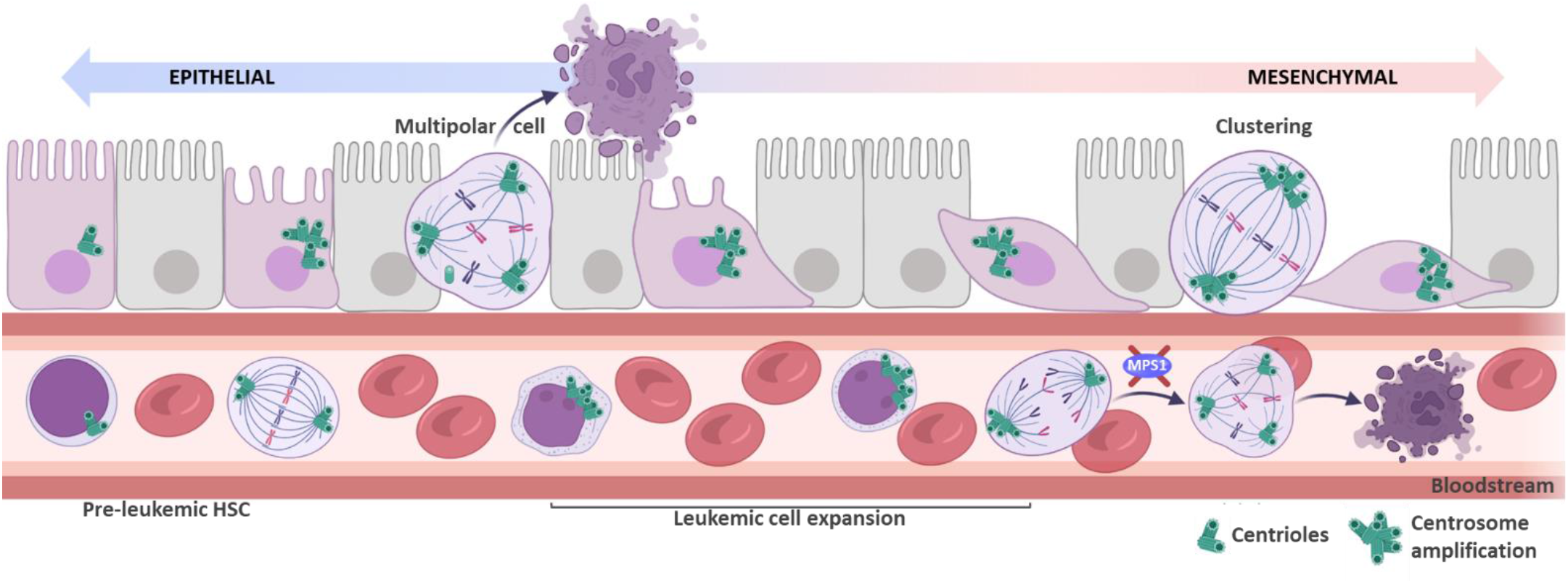

- Centrosome clustering is widespread in the NCI60 panel and particularly strong in leukemia.
- The EMT enhances the ability of cancer cells to cope with multiple centrosomes through centrosome clustering.
- Centrosome clustering gene expression peaks at the invasive stage and has prognostic value in breast cancer.
- The inhibition of MPS1 is a good strategy to promote the death of acute lymphoblastic leukemia (ALL) cells with CA.

## INTRODUCTION

Centrosomes are the major microtubule (MT) organizing centers in human dividing cells, contributing to mitotic bipolar spindle assembly and thus ensuring a balanced segregation of chromosomes. Centrosomes are composed of two microtubule-based barrel-like structures, the centrioles, and their surrounding proteinaceous material that nucleates and anchors microtubules, the pericentriolar material (PCM). The canonical centrosome duplication cycle is coupled to the chromosome cycle and tightly controls centrosome number in the cell (Gomes Pereira *et al*, 2021). However, centrosome amplification (CA), the presence of more than two centrosomes in a mitotic cell, is a hallmark of cancer and has been linked to genetic instability, invasiveness and metastatization (Chan, 2011; Godinho *et al*, 2014; Arnandis *et al*, 2018; Silkworth *et al*, 2009; Singh *et al*, 2020; Marteil *et al*, 2018; Adams *et al*, 2021). Moreover, CA was shown to trigger cancer formation in mice (Serçin *et al*, 2016; Coelho *et al*, 2015; Levine *et al*, 2017).

CA is a double-edged sword for cells. On one hand, CA can lead to the formation of multipolar mitosis triggering cell cycle arrest or cell death due to severe aneuploidy. On the other hand, centrosomes can be organized to promote pseudo-bipolar mitotic spindles, which enable cells to divide successfully (Sabat-Pośpiech *et al*, 2019). However, in this case, lagging chromosomes can be generated, which can drive chromosome instability, tumor heterogeneity, and recurrence (Ganem *et al*, 2009; Fan *et al*, 2015; Silkworth *et al*, 2009). Several cellular mechanisms allow cells to cope with CA, including the gathering or clustering of supernumerary centrosomes at the two spindle-poles (named centrosome clustering, CC) or the extrusion of extra centrioles from the spindle due to a partial (named extrusion) or complete reduction (inactivation) of their microtubule-nucleating capability, which can result from PCM loss (Gruss, 2018; Gambarotto *et al*, 2019; Sabino *et al*, 2015). The clustering of extra centrosomes during mitosis is the best-studied coping mechanism (Godinho & Pellman, 2014; Marthiens *et al*, 2012; Rhys *et al*, 2018). Previous works highlighted several cellular mechanisms important for the efficiency of CC: i) the spindle assembly checkpoint (SAC), which delays mitotic progression until all chromosomes are bioriented, ii) motor proteins associated with the mitotic spindle, such as HSET/KIFC1 or dynein (Quintyne *et al*, 2005; Chavali *et al*, 2016; Rhys *et al*, 2018) that bring centrosomes together or focus the poles, respectively and, iii) the cortical actin cytoskeleton and cell adhesion, which provide spatial cues guiding the positioning of centrosomes in two poles of the spindle via astral microtubules (Drosopoulos *et al*, 2014; Kwon *et al*, 2008; Leber *et al*, 2010; Kwon *et al*, 2015; Basto *et al*, 2008; Rhys *et al*, 2018).

It has been suggested that selective inhibition of CA coping mechanisms could provide a strong therapeutic window to specifically target cells with supernumerary centrosomes, avoiding the toxicity to normal cells and the development of resistance. Importantly, as a proof of concept, preventing CC by HSET inhibition led to multipolar mitosis and cell death in cells with CA but not in normal cells (Quintyne *et al*, 2005; Watts *et al*, 2013; Kleylein-Sohn *et al*, 2012; Li *et al*, 2015). However, the available HSET inhibitors either show a lack of efficiency and specificity or have adverse toxicity effects (Watts *et al*, 2013; Wu *et al*, 2013; Zhang *et al*, 2016a; Choe *et al*, 2018), suggesting the importance of exploring new anti-clustering drugs and/or identifying in which tissues they may be more efficient.

Despite growing interest in taking advantage of CA coping mechanisms to kill cancer cells, little is known about their prevalence in different types of tumors and their underlying molecular mechanisms. We previously used the NCI60 panel of cancer cell lines to assess the prevalence of CA in cancer cells (Marteil *et al*, 2018). As a repository of cancer diversity, the NCI60 panel contains 60 cell lines derived from nine distinct tissues, for which extensive information has been gathered along the years (gene expression, drug sensitivity amongst others), (Shoemaker, 2006; Liu *et al*, 2010; Caroli *et al*, 2020; Staunton *et al*, 2001; Zoppoli *et al*, 2012; Scherf *et al*, 2000). In this work, we take advantage of the NCI60 panel to perform the first large-scale screen to assess the frequency of CA-coping mechanisms among nine different cancer types. Our results indicate a clear correlation between CC and the loss of epithelial properties both in the NCI60 cell lines and also along breast cancer progression. Moreover, we observe that the induction of a mesenchymal state enhances the ability of breast cancer cells to cluster centrosomes. Finally, we investigated the extreme case of leukemias, which lack epithelial and cell adhesive properties, and observed the highest penetrance of CC. We further inhibited CC by targeting the SAC in leukemia, which shortens mitosis and facilitates the occurrence of multipolar mitosis and consequently, cell death. Our work suggests a new mechanistic role for the EMT transition in promoting cell viability and identifies characteristics of tumors more susceptible to drugs that interfere with CC.

## RESULTS

### Centrosome clustering is widespread in the NCI60 panel

To investigate the prevalence of CA coping mechanisms in cancer cells, we took advantage of our recent survey of centrosome abnormalities in the NCI-60 panel (Marteil *et al*, 2018). We first selected the 29 NCI-60 cell lines that were previously found to display a significant level of CA as compared to non-transformed cells (more than 12% of mitotic cells with more than 4 centrioles) (Marteil *et al*, 2018). Given the objective of systematically evaluating the prevalence of those mechanisms in each cancer type, by investigating several cell lines that are adherent and/or suspension, we opted for a screening in fixed samples (Fig. 1A). As in the original *Marteil G. et al* (2018) screen, we used two centriole distal markers, Centrin-2 and CP110, to identify bona-fide centrioles.

**Fig. 1.**
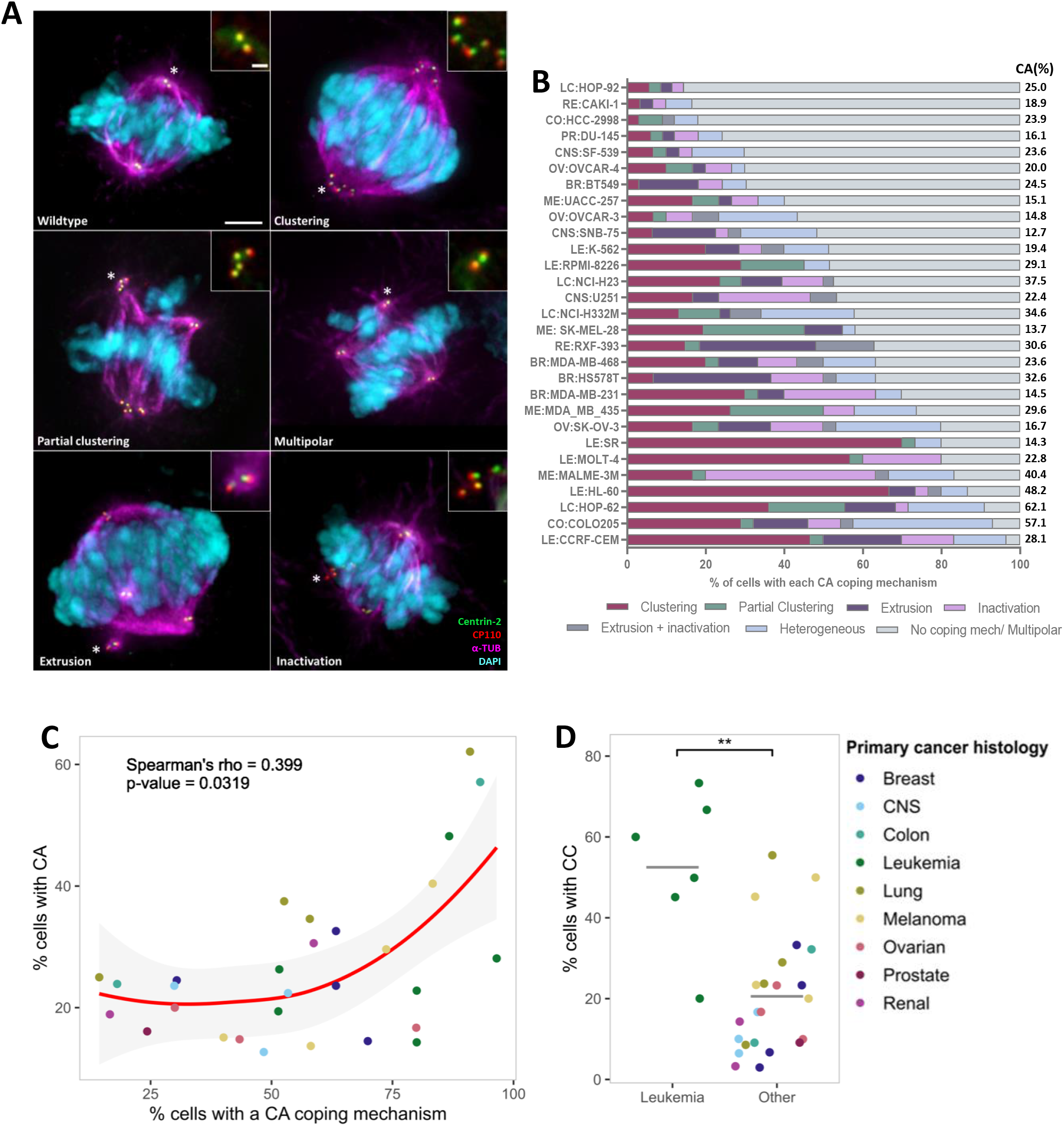
Centrosome clustering is widespread in the NCI60 panel and shows high prevalence in leukemia. **(A)** Immunofluorescence images of mitotic cells divided in six categories: **Wildtype**, cells with normal spindle with two poles and two centrioles at each pole; **Clustering**, cell with two spindle poles and at least one of them with more than two centrioles; **Partial clustering**, cell with a multipolar spindle, and at least one of the spindle poles with more than two centrioles; **Multipolar**, cell with more than two poles, each one with maximum two centrioles; **Extrusion**, mitotic spindle in which some centrioles show reduced microtubule nucleation capacity (measured by decreased α-Tubulin staining around them); **Inactivation**, mitotic cell presenting centrioles with no microtubule nucleation capacity, i.e. no clear α-Tubulin staining surrounding them. Extrusion + Inactivation, (combination of these two last mechanisms). Heterogeneous phenotypes (combinations of the different coping mechanisms) are also represented in B. Representative images show the DNA (cyan), Centrin-2 (green), CP110 (red), and the mitotic spindle (α-Tubulin -magenta). Scale bar 5 μm, insets 1 μm. α-Tubulin was removed from the insets, except for extrusion and inactivation, to facilitate visualization. **(B)** Quantification of centrosome amplification coping mechanisms in the 29 cell lines from the NCI-60 panel shown previously to display CA (Marteil *et al*, 2018). Prevalence of the different types of CA-coping mechanisms was measured as the percentage of cells with supernumerary centrioles that exhibit each mechanism. Results are depicted in the bar graph; cell lines were ranked according to the total percentage of mitotic cells showing coping mechanisms. (BR) breast, (CNS) central nervous system, (CO) colon, (LE) leukemia, (LC) lung, (PR) prostate, (RE) renal, (OV) ovary and (ME) melanoma. At least 30 mitotic cells with CA were scored per cell line. **(C)** The percentage of cells that use one or more coping mechanism(s) (X-axis) correlates with the percentage of cells that have supernumerary centrosomes (Y-axis, Spearman correlation with p-value < 0.05). Red line is the *Loess fit* (locally weighted smoothing), with 95% confidence interval in grey shaded area, added to facilitate the visualization of the correlation between CA-coping mechanisms and CC. **(D)** The percentage of cells that cluster centrosomes (CC) is significantly higher in leukemia cell lines than in other tissues. Grey lines denote the median. ^**^ p-value < 0.01 (Wilcoxon rank-sum test).

In each cell line, we investigated the presence of the following classes of coping/non coping mechanisms during mitosis: (i) clustering (CC) - bipolar spindle with at least one spindle pole containing more than 2 centrioles, (ii) partial clustering - multipolar spindle where at least one of the poles has more than 2 centrioles; (iii) extrusion – bipolar spindle in which some centrioles show decreased α-tubulin nucleation and are extruded from the spindle, not perturbing its bipolarity; (iv) Inactivation – an extreme case of extrusion in which centrioles have no clear α-tubulin nucleation; (v) extrusion + inactivation - combination of the two mechanisms and (vi) heterogeneous – bipolar spindles which exhibit combinations of the different coping mechanisms (e.g. clustering + extrusion and/or inactivation, partial clustering + extrusion and/or inactivation) and (vii) no coping mechanisms - plain multipolar spindles (with at least 3 poles with a maximum of 2 centrioles in each pole) (Fig. 1A). Our results show that the presence of coping mechanisms was higher than 40% in 22 out of 29 of the cell lines, with only three cell lines showing less than 20% prevalence of coping mechanisms (HOP-92, CAKI-1 and HCC-2998). Amongst the coping mechanisms, CC was present in all the 29 cell lines, suggesting that this mechanism is widely present in cancer cells (Fig. 1B). We also observed that CC was very heterogeneous: 12 out of the 29 cell lines exhibited less than 20% of their mitotic spindles with totally/partially clustered extra-centrosomes, 9 cell lines displayed a penetrance of between 20 and 40%, and 8 cell lines had more than 40% of mitotic spindles with clustered centrosomes. Importantly, we observed that all the 29 cancer cell lines that were screened, also presented alternative mechanisms to cope with supernumerary centrosomes. In summary, our systematic survey shows that coping mechanisms are widespread, and the percentage of cells with CA within each cell line significantly correlates with the general presence of coping mechanisms (Fig. 1B, C).

We observed that CC was particularly penetrant in leukemia (Fig. 1B, D). This is the only liquid cancer represented in the NCI60 panel, showing neither epithelial nor cell-cell adhesive properties, unlike the rest of the tissues in the panel. Recent reports show that adherent cells preserve a memory of the adhesive contacts driven by E-cadherin and established in interphase. When in mitosis, cells retain actin containing retraction fibers (RFs) linked to those sites of strong adhesion. The accumulation of RFs decreases cortical contractility in cells, which is needed to restrict centrosome movement and allow for HSET-mediated CC (Rhys *et al*, 2018; Kwon *et al*, 2008). The lack of adhesive properties in leukemia could potentially increase cortical contractility and thus explain the higher penetrance of CC observed; those properties could ideally also help predict the clustering ability of other tumors. We next set out to identify general molecular properties that correlate with clustering, in particular those related with the loss of epithelial properties.

### Centrosome clustering is associated with the loss of epithelial features and the EMT

To identify molecular properties associated with clustering, we first searched for associations between CC-ability of each cell line with its transcriptional profile, which is publicly available for 27 out of the 29 NCI-60 cell lines screened (Marteil *et al*, 2018; Reinhold *et al*, 2012). Cell lines grouped mainly by their tissue of origin, especially leukemia cell lines, which showed a very distinct transcriptional profile (Fig S1A). Given that leukemia cell lines also have a strong clustering capacity (Fig 1D), we reasoned that any correlation between their transcriptional profile and CC may partially re?ect leukemia-specific gene expression, and not necessarily be specifically associated with centrosome clustering. Additionally, we also found that the doubling time of the cell lines was inversely correlated with CC capacity (Fig S1B), and thus *per se*, could also insert a bias related to cell cycle genes in our search for molecular mechanisms specifically involved in centrosome clustering. While we cannot rule out that these two variables, tissue specific expression and doubling time, may per se affect the clustering ability, for simplification and to prevent confounding effects in our initial analysis, we opted for considering them as independent variables in our analysis. With this aim, we fitted a linear regression model that takes both effects into account as independent variables (Fig. S1C-E; see materials and methods for more details). Applying this model, we obtained a centrosome clustering signature (Table 1). We then used that signature to identify cellular pathways and components associated with centrosome clustering by performing Gene Set Enrichment Analysis (GSEA) (Subramanian *et al*, 2005). GSEA revealed a significant enrichment of centrosome-related gene sets among the positively CC-associated genes that we found (e.g., the GO-terms “Centrosome”, “Pericentriolar material”, “Centriole duplication” and “Ciliary basal body”; GSEA FDR < 0.05; Fig 2A; Fig. S2A, B, Table 2) as well as components associated with cell proliferation (e.g., G2/M checkpoint, Myc targets) that may influence centrosome clustering ability. The presence of those centrosome components may relate to specific roles in centrosome clustering that should be investigated in the future.

**Fig. 2.**
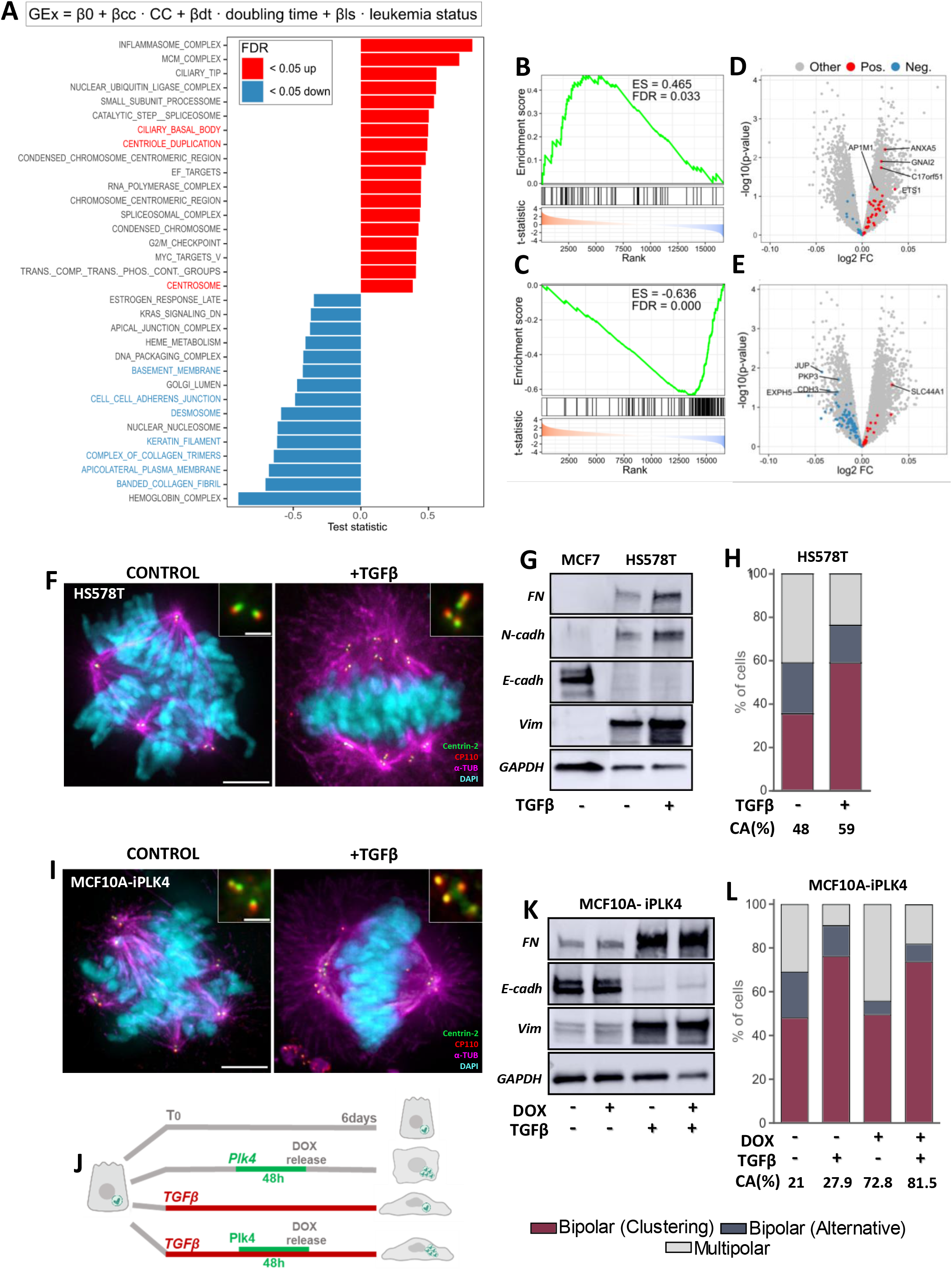
Centrosome clustering is associated with the EMT in the NCI60 panel of cancer cell lines. **(A)** Barplot showing gene sets whose down or up-regulation is significantly associated with CC (FDR < 0.05). Gene Set Enrichment Analysis (GSEA) was done on genes ranked by the t-statistic of association with CC linearly modeled as per the equation shown and described in Fig. S1D (Table 2). Ribosomal gene sets, of which several were upregulated, and general extracellular matrix gene sets, of which several were downregulated, were omitted to improve visualization and as they are frequently falsely enriched due to gene-length bias (Mandelboum *et al*, 2019). **(B, C)** GSEA results for a mesenchymal (B) (MESENCHYMAL_ROKAVEC_CCLE) and an epithelial (C) (EPITHELIAL_ROKAVEC_CCLE) gene set with genes ranked by their association with CC in the linear model (Table 3). The results indicate a positive association of mesenchymal and a negative association of epithelial genes with CC. GSEA p-values are shown. Genes are ranked by their t-statistic of association with CC and plotted on the X-axis. Genes that belong to the gene set of interest are indicated by vertical black bars. The enrichment score is plotted in green. **(D, E)** Volcano plots of differential gene expression associated with CC. Genes that belong to the mesenchymal (D) and the epithelial (E) gene sets in (B,C) are highlighted according to whether they are significantly negatively (blue) or positively (red) associated with CC. One of our top upregulated mesenchymal gene ANXA5, has been shown to promote epithelial-mesenchymal transition (EMT) in renal cancer (Tang *et al*, 2017) and JUP, the top downregulated epithelial gene found, is an important component of intercellular junctions, which help hold neighboring cells together thus inhibiting cell spreading, invasion and metastases (Aktary *et al*, 2017). **(F)** Representative images showing the effect of the treatment of HS578T cells with TGFβ (6 days). **(G)** Representative Western Blot analysis showing main EMT markers (loss of E-cadherin (E-cadh), upregulation of Fibronectin (FN), N-cadherin (N-cadh) and Vimentin (Vim)) in control cells (MCF7 -full epithelial cell line) and in HS578T cells treated with and without TGFβ for 6 days. GAPDH level was used to normalize the amount of protein loaded per lane. **(H)** Percentage of mitotic cells assemblying bipolar spindles, formed either by CC or by alternative mechanisms (Extrusion, inactivation or the combination of both), and cells showing multipolar spindles in HS578T control cells and treated with TGFβ for 6 days (A minimum of 50 cells with CA were counted per experiment (n=3)). Quantifications were performed at the end of the 6 days in both conditions. Significant p-values derived from Tukey’s multiple comparisons test performed by two-way ANOVA for changes in the % of cells undergoing clustering and multipolar spindles between control conditions and the TGFβ treatment are ^***^p= 0.0009 and ^**^p= 0.0094 respectively. The total percentage of CA is also annotated in both conditions. **(I)** Representative images showing the effect of TGFβ treatment (6 days) in MCF10A-iPlk4 cells (I). **(J)** Flowchart summarizing the experiments designed to test the effect of the EMT in CC efficiency in MCF10A-iPLK4. **(K)** Representative Western Blot analysis showing main EMT markers (loss of E-cadherin (E-cadh) and upregulation of Fibronectin (FN), and Vimentin (Vim)) in the presence or absence of TGFβ treatment conjugated with PLK4 expression induced with doxycycline in MCF10A-iPlk4 cell line. GAPDH level was used to normalize the amount of protein loaded per lane. **(L)** Percentage of mitotic cells with bipolar spindles, formed either by CC (red) or alternative mechanisms (Extrusion, inactivation or the combination of both) (blue), and multipolar spindles (grey) in MCF10A-iPLK4 control cells and treated with TGFβ for 6 days (A minimum of 50 cells with CA were counted per experiment (n = 3). CA (%) scored in MCF10A-iPLK4 in control conditions (nothing added), treated either with doxycycline or TGFβ alone or the combination of both (Doxycycline + TGFβ) are shown. Quantifications were performed at the end of the 6 days in all the conditions. Note that the level of CA is higher than usual in basal conditions probably caused over time by leaky Plk4-expression in the absence of doxycycline. Significant P-values derived from Tukey’s multiple comparisons test performed by two-way ANOVA are the following: **Clustering**: ^****^p< 0.0001 for Control vs. TGFβ; ^***^p= 0.0002 for Control vs. TGFβ-Dox; ^***^p= 0.0001 for Control-Dox vs. TGFβ, ^***^p= 0.0005 for Control-Dox vs. TGFβ-Dox; ^***^p ≤ 0.0005. **Multipolar**: ^**^p=0.0019 for Control vs. TGFβ; ^****^p<0.0001 for Control-Dox vs. TGFβ; ^***^p=0.0002 for Control-Dox vs. TGFβ-Dox. **Alternative mechanisms**: ^*^p=0.0386 Control vs. Control-Dox. Images show the DNA in cyan, centrioles in green (Centrin-2) and red (CP110), and the spindle in magenta (α-Tubulin). Scale bar 5 μm, insets 1 μm.

Importantly, we observed an association between CC-capacity and EMT-related events (GSEA FDR < 0.05; Fig. 2A, blue bars. Table 2), such as the loss of epithelial properties; including downregulation of cell-cell adhesion factors (adherens junctions or desmosomes), factors responsible for maintaining cell polarity or involved in the contact with the basement membrane (Roche, 2018). To further investigate the correlation between the EMT and CC, we performed GSEA with additional gene sets: i) a curated list of custom gene sets for epithelial and mesenchymal factors obtained from Mak *et al*, 2016 and Rokavec *et al*, 2017 (named Rokavec signatures from TCGA, Rokavec signatures from CCLE, and Mak signatures from TCGA), ii) reduced signature based on the literature (named “Epithelial curated” and “Mesenchymal curated”, Table 3) (Lee *et al*, 2006; Lee & Nelson, 2012; Gröger *et al*, 2012; Zhao *et al*, 2015), and iii) a combination of epithelial or mesenchymal genes from all signatures (“Epithelial total”, “Mesenchymal total”). Our results show that epithelial signatures are strongly enriched among the negatively CC-associated genes (Fig. 2C,E; S2C, Table 3). One of the most striking features of the invasive breast cancer is the loss of expression of the intercellular adhesion molecule E-cadherin, recently shown as a prerequisite for cells to cluster centrosomes (Corso *et al*, 2020; Padmanaban *et al*, 2019; Rhys *et al*, 2018). This loss of expression was found consistently in cell lines with efficient CC in the NCI-60 panel (Fig. S2D, Table 1). In fact, one of the top proteins negatively associated with CC was PDLIM1 (Fig S2D, Table 1), whose downregulation has been associated with metastatic potential in colorectal cancer through destabilization of the E-cadherin/β-catenin complex (Chen *et al*, 2016). Most mesenchymal gene sets tested were enriched among the genes positively associated with CC (Fig. 2B,D; S2C, Table 3), highlighting the importance of loss of epithelial properties, as well as of the acquisition of mesenchymal features for efficient CC.

### The EMT enhances the ability of breast cancer cells to cope with multiple centrosomes

Our results led us to hypothesize that the loss of epithelial properties might facilitate the clustering of extra-centrosomes in cancer cells. CA is often observed in breast cancer and underlies tumor aggressiveness and prognostic of worse outcomes (Denu *et al*, 2016; Singh *et al*, 2020; Lingle *et al*, 2002; Salisbury; Zhang *et al*, 2020; D’Assoro *et al*, 2002; Pannu *et al*, 2015a; Marteil *et al*, 2018). Metastasis of breast cancer cells requires decreased cell–cell adhesion and cells to undergo the EMT, in which epithelial cells acquire a motile mesenchymal phenotype (Roche, 2018). We therefore proceeded to validate our hypothesis in basal breast cancer. We identified breast cancer cell lines with varying ability to cluster in our experimental conditions: amongst the basal cell lines investigated, two (MDA-MB-231, MDA-MB-468) displayed major levels of clustering (33.3 and 23.3%, respectively), while the cell lines HS578T and BT549 were less efficient at clustering (6.7 and 3%, respectively) (Fig. 1B-BR, 1D). A recent report showed that HS578T and BT549, despite showing loss of E-cadherin (Tryndyak *et al*, 2010), are in an intermediate state of EMT (Margaron *et al*, 2019), which may explain their compromised ability to cluster centrioles. We chose HS578T for the analysis, as BT549 cell line frequently shows acentrosomal poles (Kleylein-Sohn *et al*, 2012). Our hypothesis being true, the induction of a full mesenchymal phenotype in HS578T should increase its ability to cluster centrioles. Therefore, we exposed this cell line to TGFβ for 6 days to allow cells to acquire a full mesenchymal phenotype as previously reported (Lindley & Briegel, 2010) (Fig. 2F-H).

The effect of TGFβ was first validated by western blot (WB) (Fig. 2G). A hallmark of EMT is the upregulation of N-cadherin followed by the downregulation of E-cadherin (Loh *et al*, 2019). As control, we analysed EMT makers in the epithelial breast cancer cell line MCF7, which shows high levels of E-cadherin and full absence of the mesenchymal markers Fibronectin, N-cadherin and Vimentin, consistent with an epithelial phenotype. The previously described intermediate EMT state of the HS578T cell line explains the presence of basal levels of the mesenchymal markers Fibronectin, N-cadherin and Vimentin and total absence of E-cadherin (Margaron *et al*, 2019). However, the observed increase in fibronectin, N-Cadherin, and vimentin, 6-day after exposure to TGFβ, was consistent with an increased mesenchymal phenotype as previously reported (Dongre *et al*, 2017). As predicted, the percentage of cells with CA undergoing bipolar mitosis due to centrosome clustering significantly increased upon TGFβ treatment in detriment of multipolar mitoses (Fig. 2F, H).

Cancer cells show a variety of different uncharacterized mutations that may influence our results. To ask more directly whether mesenchymal characteristics could favour the survival of cells with extra centrosomes, we induced CA, concomitantly with the EMT, in normal breast cells. For this, we used a doxycycline-inducible PLK4 epithelial cell line MCF10A (Denu *et al*, 2020; Arnandis *et al*, 2018) treated with TGFβ to promote EMT (Fig. 2I-L) according to the protocol shown in Fig. 2J.

We first confirmed the acquisition of a mesenchymal phenotype by WB and observed a clear downregulation of E-cadherin and upregulation of N-cadherin, Vimentin and Fibronectin protein levels (Fig. 2K). TGFβ treatment did not induce a significant increase in CA in MCF10A cells (Fig. 2L). However, as expected, CA (>4 centrioles per mitotic cell) increased from 25 to 70% upon PLK4 overexpression with doxycycline (Fig. 2L). We then assessed the centrosome clustering ability of MCF10A-iPLK4 cells with and without incubation with TGFβ. Induction of a mesenchymal phenotype after CA-induction significantly increased CC and reduced the percentage of multipolar spindles (Fig. 2L). We also observed a consistent increase of centrosome amplification throughout the different experiment replicates in HS578T and MCF10A-iPLK4 cell lines when exposed to TGFβ, suggesting the cells with CA have acquired survival advantages (Fig 2 H, L). In summary, our results support the idea that the EMT promotes the survival and thus the maintenance of cells with CA within the population. Consequently, we would expect CC to be stronger as breast tumors go through the EMT.

### Expression levels of centrosome duplication and clustering genes concomitantly change along breast cancer malignant progression and peak at the invasive stage

Clustering is arduous to analyze accurately *in situ* in tumors due to the difficulty of identifying mitotic cells and observing the whole mitotic spindle and all centrioles within the histochemistry cut (Morretton *et al*, 2022). As a proxy to investigate at which stage CA and CC occur *in vivo* during breast tumorigenesis, we followed the expression profile of centriole duplication and clustering related genes (Table 03) throughout the several epithelial proliferative breast diseases that are considered as distinct conditions of breast cancer progression (normal breast epithelium (N), simple hyperplasia (SH), atypical hyperplasia (AH), ductal carcinoma in situ (DCIS) and invasive ductal carcinoma (IDC)) (Fig. 3A, Fig S3A-E) (Simpson *et al*, 2005; Bombonati & Sgroi, 2011; Hartmann *et al*, 2014). We first mined and compiled transcription profiles from the different progression stages of breast cancer from public databases (Fig S3A-E, Table 04). We found that centriole duplication genes start to be upregulated in AH and their expression increases further throughout DCIS and IDC (Fig. 3B, D; S3D). This early upregulation of centriole duplication genes is supported by the fact that CA was previously found in pre-cancerous and pre-invasive breast carcinomas (Pihan *et al*; Pannu *et al*, 2015a; Guo *et al*, 2007; Zhang *et al*, 2020). Interestingly, the expression of genes which are required for CC was only upregulated in the most aggressive stages of breast cancer progression, particularly IDC (Fig. 3C, E, Fig. S3E), precisely when EMT takes place and cells become more mesenchymal and able to metastasize. For instance, while a major trigger of centriole formation, PLK4, was up-regulated already in atypical hyperplasia (AH) and kept its high expression along malignant progression (Fig. S3F), a major regulator of CC, KIFC1/HSET only started to be significantly up-regulated later, from DCIS onwards (Fig. S3G). These results suggest that the ability to cluster centrosomes might allow for cells with CA to survive better in malignant stages and therefore potentially contribute to worst cancer outcomes.

**Fig. 3.**
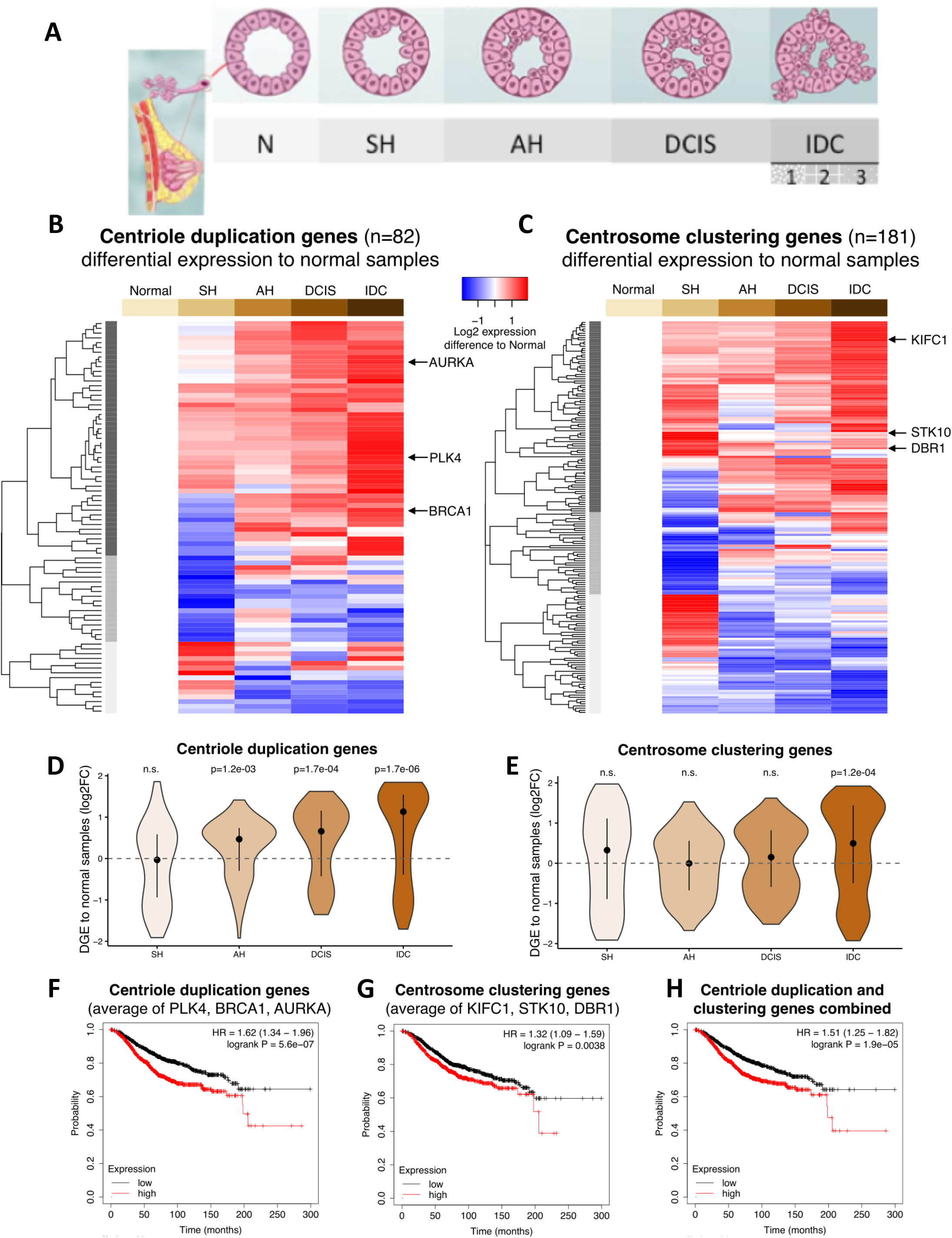
Evidence for *in vivo* association of mesenchymal characteristics with centriole amplification and clustering. **(A**) A breast cancer progression model was used to retrieve data for the different stages of disease development (Table 4). These include: N, normal; SH, simple hyperplasia, AH, atypical hyperplasia; DCIS, ductal carcinoma *in-situ*; IDC, invasive ductal carcinoma. Elston Grading (Elston & Ellis, 1991) was used to stage the three IDC stages. **(B, C)** mRNA expression profiles at the different breast cancer stages of **(B)** centriole duplication genes (manually curated) and **(C)** centrosome clustering genes (compiled from 3 RNAi-screens: ((Drosopoulos *et al*, 2014; Leber *et al*, 2010; Kwon *et al*, 2008), Table 4). Genes were selected for display if differentially expressed against normal breast tissue in at least one comparison (FDR < 0.05) and ordered based on unsupervised hierarchical clustering (the 3 main gene clusters are highlighted with different shades of gray). Three representative genes involved in centriole duplication and in centrosome clustering and upregulated during breast cancer progression are highlighted in (B) and (C) respectively. The displayed log2 expression values are the median for the samples of each stage and were centered to the Normal (N) breast median (i.e., they are the log2 fold-changes of the median when compared to Normal breast samples). **(D, E)** Violin plots showing the distribution of differential gene expression (DGE) values (log2 fold-change (log2FC) of median) of **(D)** centriole duplication and **(E)** centrosome clustering genes at the different breast cancer stages against normal breast samples. Black points and lines represent the median +/-upper/lower quartiles. P-values are from one-sample Wilcoxon tests against 0; n.s. non-significant (p > 0.05). **(F-H)** Kaplan-Meier plots of disease-specific survival analysis of breast cancer (> 2000 METABRIC samples), based on the combined expression of **(F)** centriole duplication genes *PLK4, BRCA1* and *AURKA*, **(G)** centrosome clustering genes *KIFC1, STK10* and *DBR1* and **(H)** the six genes altogether, show that their up-regulation is significantly associated with worse prognosis. Each combined expression score was calculated as the sum of the log2-transformed and median-centered expression values of the considered genes.

Our data suggests that CC is associated with worse prognosis. To further test this idea, we used gene expression and clinical data from multiple breast cancer datasets (see Material and Methods; (Győrffy, 2021)) and assessed the combined effect of the upregulation of both known centrosome components involved in centriole duplication, function and tumorigenesis (PLK4, BRCA1 and AURKA, (Qi *et al*, 2022; Magnaghi-Jaulin *et al*, 2019; Tkach *et al*, 2022; Moyer & Holland, 2019)), and regulators of CC. For this purpose, we chose the well-characterized regulator of centrosome clustering, KIFC1 (Pannu *et al*, 2015b; Vitre *et al*, 2020; Chavali *et al*, 2016), one suggested regulator of centrosome clustering and also present in our signature, DBR1 (Table 1, (Kwon *et al*, 2008) and a gene that was upregulated in the cell lines that cluster better, STK10 (Table 1). The up-regulation of those genes, both independently and combined, was significantly associated with worse prognosis (Fig. 3F-H).

Our results show that the expression levels of centrosome duplication and clustering genes are concomitantly increased along breast cancer malignant progression and peak at the invasive stage. Our data also suggests that breast cancer tumors with more mesenchymal properties should be better candidates for therapies targeting CC.

### Centrosome clustering is particularly strong in leukemia

Therapies focused on inhibiting CC have not been particularly successful, in part as they were targeting tissues where clustering was not very penetrant (Watts *et al*, 2013; Wu *et al*, 2013; Zhang *et al*, 2016a). We wondered whether targeting tissues which show a higher clustering ability could allow a more efficient treatment. Given that five of the six leukemia cell lines present in the NCI60 panel had over 40% ability to cluster (Fig. 1B), our results suggest that hematological malignancies are particularly capable of clustering their supernumerary centrosomes (Mann Whitney U p-value < 0.01; Fig. 1D) and could positively respond to CC-targeting therapies. Within the panel, three out of six leukemia cell lines are acute leukemia (CCRF-CEM, MOLT-4 and HL-60) and among them, two are from the lymphocytic lineage (CCRF-CEM and MOLT-4), i.e., acute lymphoblastic leukemia (ALL). The latter is a type of leukemia mostly diagnosed in children and adolescents and responsible for a third of childhood cancer deaths. While more than 80% of children with ALL survive following contemporary therapies (Pui *et al*, 2015; Hunger *et al*, 2012), the 5-year overall survival (OS) rate dramatically drops to ∼50% after the first relapse (Oskarsson *et al*, 2016; Rheingold *et al*, 2019). The OS rates for adult patients with ALL are lower than 50% % (Patel *et al*, 2020) and therefore there is an urgent need for implementation of new strategies. Lower survival rates in ALL often include TP53 mutations enrichment at relapse (Richter-Pechańska *et al*, 2017; Liu *et al*, 2017) or a history of previous chemotherapy treatments, both being previously associated with CA (Fukasawa *et al*, 1996; Demoor-Goldschmidt & de Vathaire, 2019; Rheingold *et al*, 2003; Varan & Kebudi, 2011; Lorenzi *et al*, 2011; Oeffinger *et al*, 2006). Given that toxicity associated with chemotherapy and refractory relapse significantly compromise overall survival of ALL patients, we further investigated whether CC could be a target in treating this type of leukemia.

We first validated our initial screening results, by analyzing the CA and CC levels in 4 additional ALL cell lines outside of our panel: HPB-ALL, JURKAT, DND41 and P12 (Fig. 4A-C). Three of the cell lines, JURKAT, DND41 and P12, showed more than 60% of CA, while HPB-ALL displayed 26% of CA, suggesting that CA is a general feature of ALL. In addition, more than 50% of the cells with CA in all cell lines showed clustered centrioles (Fig. 4C), further supporting the idea that ALL cells show efficient CC. To further validate our results *in vivo*, we used six Patient-Derived tumor Xenograft (PdX) models of ALL. ALL PdXs are the standard for preclinical research, able to recreate the tumor microenvironment and accurately replicate growth and diversity of tumor cells and tumor progression (Woiterski *et al*, 2013; Wang *et al*, 2017; Richter-Pechańska *et al*, 2018). As experimental controls, we used two normal thymocyte subsets obtained from pediatric thymic biopsies, which constitute the ALL-normal counterpart, as ALL can result from the transformation of thymic cell precursors (Cordo’ *et al*, 2021). In the six PdXs analyzed, we observed between 15% and 25% of the interphasic cells with CA and between 9 and 25% of CA in mitotic cells (Fig 4D). As the majority of thymocytes are not dividing, mitotic Fig.s were scarce, leading us to score CA in interphasic cells. We counted less than 6% of CA in the two thymocyte populations analyzed, which suggests that CA is a specific feature of leukemia. Importantly, more than 70% of mitotic cells with supernumerary centrosomes in the ALL PDXs showed CC (Fig 4E, F). We conclude that CC is highly penetrant in ALL and likely in leukemia in general.

**Fig. 4.**
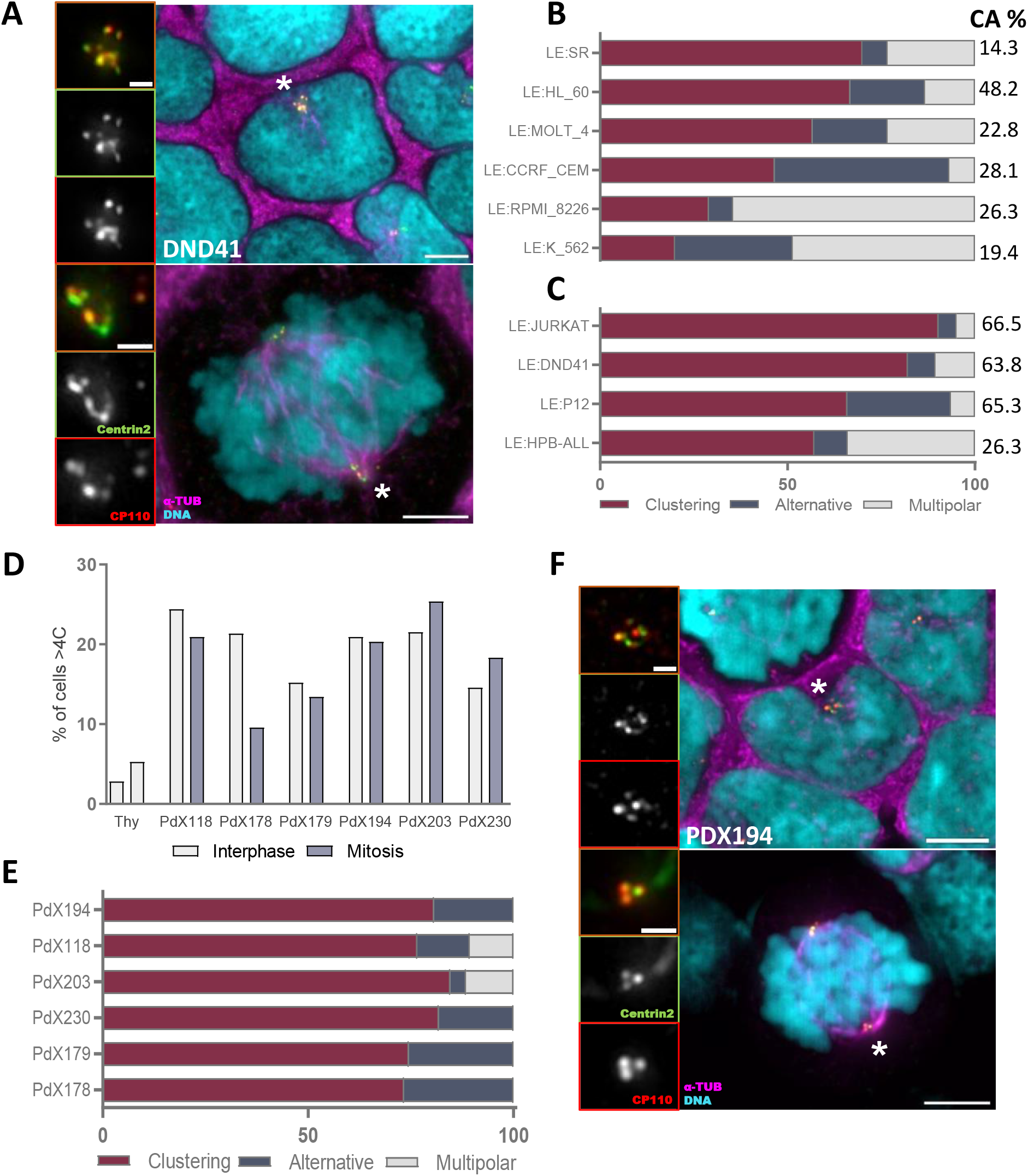
Centrosome clustering is particularly strong in leukemia cells. **(A)** Representative immunofluorescence images of interphasic (top) and clustered (bottom) mitotic DND41 cells. **(B**,**C)** Quantification of the percentage of centrosome clustering (red), alternative CA coping mechanisms (centrosome extrusion, inactivation or the combination of both (blue)), and multipolar spindles (mitotic cells with more than 2 poles (grey)) scored in (B) the NCI60 leukemia cell lines and **(C)** four alternative T-ALL cell lines. On the right column, the percentage of CA in each cell line is shown. N ≥ 100 mitotic cells with CA were analyzed in each cell line. **(D)** Comparison of the percentage of interphasic and mitotic cells with CA (>4 centrioles) in the T-ALL PdXs analyzed, as well as in two thymocyte subsets of cells derived from two distinct patients in interphase (no mitotic cells could be found). More than 100 cells were analyzed in each sample. **(E)** CA coping mechanisms present in 6 human T-ALL PdXs were scored as in (B,C). N ≥ 100 mitotic cells with CA were analized in each cell line. **(F)** Representative immunofluorescence images of PdX194-derived interphasic (top) and clustered (bottom) mitotic cells. For all immunofluorescence images, the DNA is shown in cyan, centrioles in green (Centrin-2) and red (CP110) and the spindle in magenta (α-Tubulin). White asterisks depict the area or pole magnified in the respective inset. Scale bar 5 μm, insets 1 μm

### MPS1 inhibitors kill Acute Lymphoblastic Leukemia (ALL) cells by preventing CC and promoting multipolar spindles

Clinical targeting of cell cycle and mitotic kinases in solid and hematological malignances, such as Aurora or Polo-like kinases inhibitors, have stalled in late-stage clinical trials due to lack of efficacy or significant adverse effects (Olmos *et al*, 2011; O’Connor *et al*, 2019).

Targeting CC might be a promising therapeutic target for leukemia patients. Several spindle assembly checkpoint components, motor proteins and MT and actin cytoskeleton related molecules have been proposed to be useful in the targeting of clustering mechanisms (Sabat-Pośpiech *et al*, 2019). Interestingly, spindle assembly checkpoint components, such as MPS1, are overexpressed in a variety of tumor types (Salvatore *et al*, 2007; Landi *et al*, 2008; Daniel *et al*, 2011; Miao *et al*, 2014; Zhang *et al*, 2016b; Ling *et al*, 2014; Tannous *et al*, 2013) and correlate with recurrence and poor survival (Xie *et al*, 2016). In agreement, the inhibition of MPS1 in mouse xenograft models of human cancer cell lines showed to be efficient without toxicity associated (Libouban *et al*, 2017; Simon Serrano *et al*, 2020; Maia *et al*, 2015; Jemaà *et al*, 2013; Tardif *et al*, 2011; Kusakabe *et al*, 2015b; Wengner *et al*, 2016; Colombo *et al*, 2010). Moreover, recent reports have also shown the preclinical effect of MPS1 inhibitor in ALL cell lines and also the inhibition of colony formation in a B-ALL patient (Kaci *et al*, 2019; Libouban *et al*, 2017). Therefore, we hypothesized that leukemia cells with CA and CC, in contrast to normal cells, could have increased dependency on the spindle assembly checkpoint to ensure proper clustering and bipolar formation, thus explaining why MPS1 inhibitors may have an advantage over other targeted anti-proliferative therapies.

We tested whether MPS1 inhibitors could kill ALL cells through targeting CC. First, we assessed the effect on cell viability of 5μM of the highly selective MPS1 inhibitor AZ3146 during 72h in four ALL cell lines (DND41, JURKAT, HPB-ALL and P12) (Fig. 5A-B), a similar dosing and time as published before (Jin *et al*, 2020). We saw a massive decrease in cell viability in most ALL cell lines analyzed, except for HPB-ALL, which has a longer doubling time (more than 50 hours) and may thus need treatment for longer periods of time (Fig 5B). To further investigate whether the effect of MPS1-inhibition is linked to CA, we compared cell viability after AZ3146-treatment of cancer cells without CA (HeLa), a non-cancerous cell line (hTert RPE-1) and two ALL cell lines with high level of CA (JURKAT and DND41) (Fig. 5C). AZ3146 treatment strongly decreased ALL cell viability in a dose-dependent manner. While HeLa viability was also compromised, which may result from its well characterised aneuploidy (Callaway, 2013), the effect was significantly milder as compared to ALL cells. Importantly, the effect of the MPS1 inhibitor was almost innocuous for the non-tumoral cell line RPE, supporting previous reports (Maia *et al*, 2015). Additionally, decrease in viability in ALL was, at least in part, a consequence of the inhibition of CC, as MPS1 inhibition markedly increased the percentage of cells forming multipolar spindles (Fig. 5D) (52.3, 51.1, 18.8 and 58.7% of increase of multipolar cells in DND41, JURKAT and HPB-ALL and P12 respectively). Our results suggest that MPS1 is a promising target to treat cancer cells, specifically for tumor types with high degree of CA such as ALL, where the MPS1 inhibitor promotes cell death by increasing the frequency of multipolar mitoses.

**Fig. 5.**
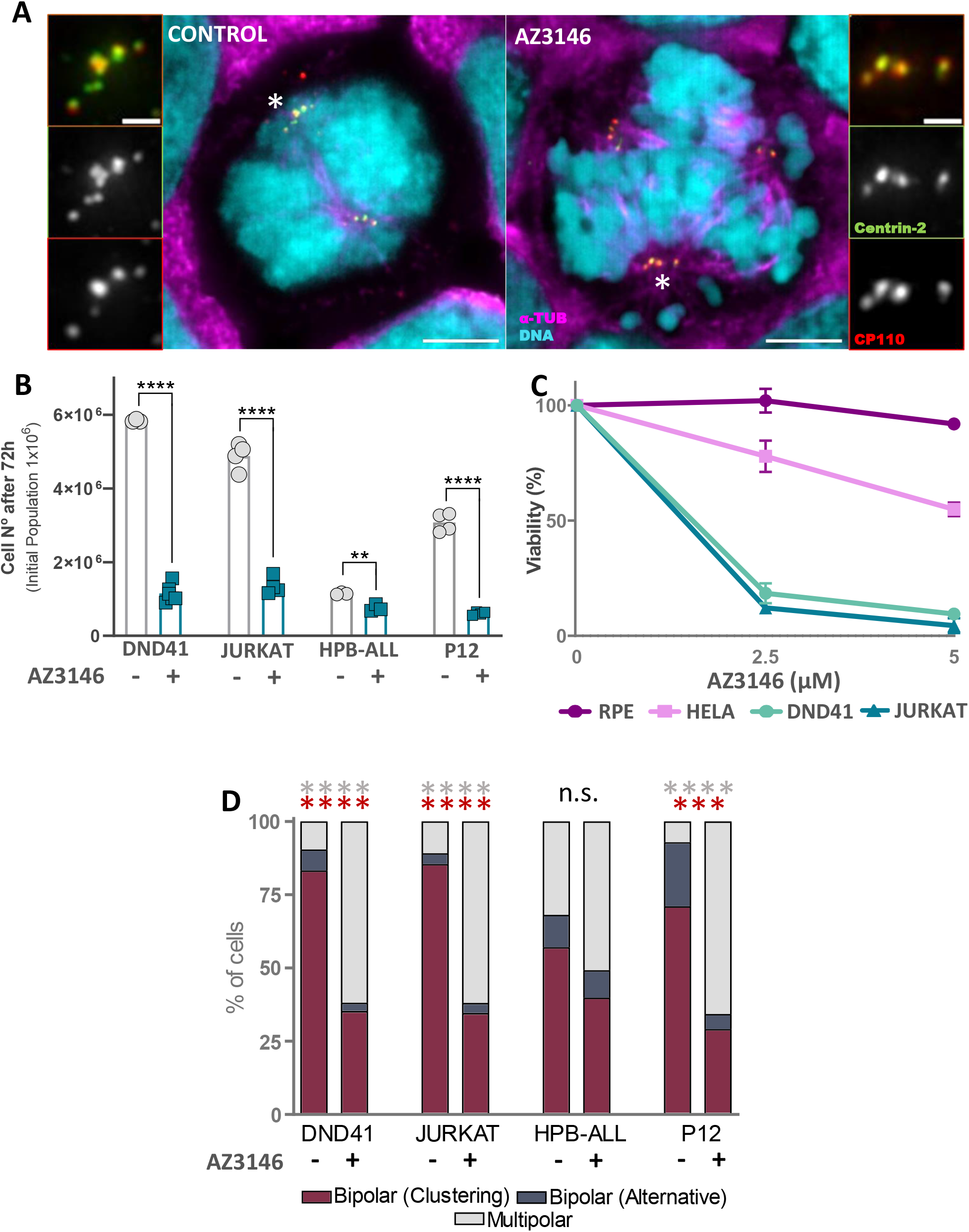
The MPS1 inhibitor AZ3146 selectively kills ALL cells by promoting multipolar spindles. **(A)** Representative immunofluorescence images of bipolar and multipolar spindles of DND41 cells treated with DMSO (left) or with 5μM of AZ3146 (right). DNA is shown in cyan, centrioles in green (Centrin-2) and red (CP110) and the spindle in magenta (α-Tubulin). White asterisks indicate the pole magnified in the respective inset. Scale bar 5 μm, insets 1 μm. **(B)** Quantification of cell survival in four ALL cell lines (DND41, JURKAT, HPB-ALL, P12) in the presence of 5μM of AZ3146. Vehicle treatment (DMSO) was used as control condition. 1×10 cells were seeded at T0 and the number of viable cells was quantified 72h later. N=2 experiments. **(C)** MTT viability assay showing the differential cytotoxic potential of AZ3146 over two ALL cell lines (JURKAT, DND41), the cancerous HeLa and the non-tumoral hTertRPE-1 cell lines. Data represent the percentage of viable cells in the presence of different AZ3146 doses (2.5μM and 5μM) upon 72h treatment. N=2 experiments. Significant p-values derived from Tukey’s multiple comparisons test performed by one-way ANOVA to evaluate AZ3146 treatment toxicity are the following: For the AZ3146 concentration of 2.5 μM: ^**^p=0.076 RPE vs HELA; ^****^p<0.0001 RPE vs. JURKAT and DND41; ^****^p<0.0001 HELA vs. JURKAT and DND41. For the AZ3146 concentration of 5 μM: ^****^p<0.0001 RPE vs. HELA; ^****^p<0.0001 RPE vs. JURKAT and DND41 and ^****^p<0.0001 HELA vs. JURKAT and DND41. Changes in viability between JURKAT and DND41 are not significant (p>0.05) at any of the concentrations. **(D)** Analysis of the percentage of bipolar (clustering or alternative mechanisms) and multipolar spindles of mitotic cells with CA after 72h of treatment with AZ3146 (5μM). More than 50 mitotic cells were analyzed per experiment. N=2. The colors of the asterisks refer to the CC coping mechanism compared in each pair of treatments per cell type. *For graphs B and D, p*-values are derived from unpaired two-tailed *t*-test. ^**^p ≤ 0.01; ^***^p ≤ 0.001; ^****^p ≤ 0.0001.

## DISCUSSION

Using the NCI60 panel of cell lines from 9 different tumor types, we previously showed that CA is a hallmark of cancer (Marteil *et al*, 2018). Here we investigated the mechanisms those cancer cells use to cope with CA and unveiled CC as the major mechanism used by cells to make pseudo-bipolar mitotic spindles and produce viable progeny. We then investigated the molecular mechanisms promoting CC in cancer and report a new role for the loss of epithelial properties and the EMT in that process. In agreement with that finding, we observed the strong clustering ability of leukemia, which are non-adherent cells and thus devoid of epithelial properties. Finally, we tested whether targeting clustering by inhibition of the spindle assembly checkpoint could be used as a therapeutic strategy in acute lymphoblastic leukemia (ALL), and showed that this treatment prevents CC and promotes cell death of the cancer cells.

While we mostly focused on CC, our survey provides novel information on the frequency of alternative mechanisms to cope with CA in nine different cancer tissues, opening new avenues for further research. For example, we often found examples of centrosome extrusion, defined as an apparent decrease in centrosome microtubule nucleation capacity and consequent exclusion from the spindle. A similar phenotype was previously reported in BT-549, a breast cancer cell line, where supernumerary active centrosomes were shown to be transiently displaced from the mitotic spindle during metaphase until anaphase transition, allowing for bipolar spindle formation (Kleylein-Sohn *et al*, 2012). Other alternative mechanism found in the cancer cell lines is the inactivation of extra centrosomes, allowing only two centrosomes to function as the microtubule organizing centers during mitosis. Centrosome inactivation has previously been described in fly neuroblasts and is characterized by a decrease in PCM content and consequently, in microtubule nucleation capacity (Basto *et al*, 2008). It is known that the MTOC activity of the centrosomes is regulated not only by the accumulation of PCM but also by their phosphorylation by mitotic kinases; however, the mechanisms leading to selective removal or inactivation of centrosomes remain a mystery. Future integration of the prevalence of each coping mechanism found in our screen with the available information on the NCI-60 gene expression and drug data may provide novel insights into their underlying molecular mechanisms and also shed light into the discovery of new drugs to selectively target those mechanisms in specific tumor types.

CA has been observed early in tumor formation (Pihan *et al*, 2003; Lopes *et al*, 2018) and is sufficient to initiate spontaneous tumorigenesis in several tissues in mice (Levine *et al*, 2017). However, it was unclear how cells could tolerate CA as it leads to multipolar mitosis and death. Here we observed that the EMT promotes adaptation to multiple centrosomes through clustering, providing a direct mechanistic link between two common characteristics of tumors with poor prognosis (mesenchymal state and CA). This goes in line with a recent report where it was shown that the loss of E-cadherin, which is commonly observed in cancer-related EMT, provides tolerance to CA due to the improvement of CC (Rhys *et al*, 2018). Given that it was shown that CA in patient-derived tumor cells reduces over time with culturing (Mittal *et al*, 2017) it is possible that *in vivo*, the activation of signalling pathways involved in EMT could ensure cell adaptation, survival and maintenance of cells with multiple centrosomes in the population. Several studies have associated CA with early events of the EMT (Rhys *et al*, 2018; LoMastro & Holland, 2019). Recent reports suggested that the presence of CA disrupts cytoarchitecture towards a more mesenchymal phenotype, also increasing their invasive properties (Arnandis *et al*, 2018; Godinho *et al*, 2014; Adams *et al*, 2021). Importantly, CA was able to drive spontaneous mesenchymal sarcomas and lymphomas *in vivo* unlike epithelial tissues such as the skin, the liver or the lungs (Levine *et al*, 2017). Overall, although the role of CA in changing cell characteristics towards a more mesenchymal phenotype has been already explored, our results point to a possible feedback loop where the EMT also contributes to the survival of cells with CA. A deeper understanding of the synergistic effects of the presence of CA and the EMT program might lead to the development of improved therapeutic strategies in epithelial tumors.

A gene expression signature for CA has been proposed, in which the increased cumulative expression of specific genes was associated with worse prognosis in a breast cancer cohort as well as with tumor incidence and progression in a pan-cancer analysis (de Almeida *et al*, 2019; Ogden *et al*, 2017). Here we determined that the up-regulation of not only centriole duplication genes but also CC genes, both independently and combined, are significantly associated with worse patient prognosis (Fig. 3). Therefore, we provide a useful tool that may be particularly relevant in the clinic to assess tumor malignancy and the possibility of recurrence. Further, it may also help in the election of more personalized treatments for patients.

Although radiotherapy and chemotherapy are currently effective methods to treat leukemia, refractory disease, relapse and toxicity remain challenging in part due to the complex and poorly understood patterns of toxicity and side effects associated (Marks *et al*, 2009). Moreover, despite the clonal heterogeneity of ALL, which limits current targeted therapy options for patients, we identify CC as a mechanism that the majority of ALL cells with CA use to survive, becoming an attractive targeting strategy. Our work also allowed us to uncover the importance of the spindle assembly checkpoint (SAC) to maintain leukemia cell viability, as inhibition of MPS1 significantly increased multipolar divisions and cancer cell death (Fig. 5). Given that the inhibition of MPS1 was significantly more capable of killing cells with CA, this approach might be particularly promising as we expect it to be effective also for other tissues with high level of CA and CC. In fact, combination therapy with chemotherapeutic drugs in triple negative breast cancer (TNBC) have already entered in phase 1 clinical trials which set the potential of the MPS1 inhibition (NCT03328494; NCT05251714).

Another potential advantage of the inhibition of the SAC could be the treatment of metastatic TNBC, since circulating tumor cells have high levels of CA and show good clustering capacity (Singh *et al*, 2020). It is clear by now that CC is an essential mechanism for chromosome instability, which contributes, between other malignant features such as drug resistance, to a high risk of tumor recurrence (Ganem *et al*, 2009; Holland & Cleveland, 2009; Krämer *et al*, 2011; Pannu *et al*, 2015b; Fan *et al*, 2021). Targeting CC might prevent the major cause of mortality in leukemia patients after infection, which is their relapse (Kiem Hao *et al*, 2020).

One limitation of our study on the CA coping mechanisms is that we include a relatively small cohort of cancer cells, as only 28 out of 60 cell lines were found to have a significant percentage of CA within the NCI60 panel. However, we managed to have a representation of all the tissues in that panel, which gives an idea on how efficient cells within a tissue are to cope with CA. We also envision limited efficacy of the single treatment with the MPS1 inhibitor AZ3146 depending on the tumor type, as it was previously described for other MPS1 inhibitors (Kusakabe *et al*, 2015a; Tardif *et al*, 2011; Tannous *et al*, 2013; Martinez *et al*, 2015). Nonetheless, our findings suggest that low levels of CA and/or CC in those tumor types may explain the current need of combination therapies to also target cells without CA (Maia *et al*, 2015, 2018). Therefore, the analysis of the level of CA and CC in tumors before further application of single or combination therapies with MPS1 inhibitors could improve dosage determination, drug toxicity and thus determine the success of the treatment. The role of the loss of E-cadherin in improving CC has been attributed to an increased cell contractility due to the loss of mechanical cues with adjacent cells. This restricts centrosome movement, bringing them in close proximity so that the kinesin HSET/KIFC1 can promote MTs cross-linking and the subsequent centrosome clustering (Kwon *et al*, 2008). As leukemia cells have an increased contractility due to the lack of cell adhesions, treating cells with agents that increase their adhesive properties, for example by modulating E-cadherin expression, could inhibit CC and promote cell death. Indeed, restoration of E-cadherin expression and E-cadherin-mediated cell adhesion in leukemia inhibited growth and colony formation (Rao *et al*, 2011). Further insights into the functional interplay between the inhibition of the SAC and the clustering machinery hold promise for design of smart innovative combinatorial approaches to personalized treatment of cancer.

## MATERIAL AND METHODS

### Primary samples, cell lines and culture

#### NCI60 cell lines

Cell lines were cultured in their respective media (see Table 5) supplemented with 10% Fetal Bovine Serum (FBS, Biowest) and PSA, an antibiotic (100 U/mL Penicillin and 0.1 mg/mL Streptomycin) and antimycotic (0.25 μg/mL Amphotericin B) solution from Sigma. Cell lines were maintained at 37°C with 5% CO2 atmosphere. Specific supplements are indicated in Table 5.

#### Additional cell lines

JURKAT, DND41, HBT-ALL and P12 cells were cultured in RPMI 1640. HeLa cells in DMEM and RPE1 cells in DMEM/F-12 (Gibco) at 37 °C and 5% CO_2_. All media were supplemented with 10% FBS, 1% PSA and 2mM L-glutamine. MCF10A-iPLK4 cells (obtained from S. Godinho’s laboratory) were grown in DMEM/F12 (Invitrogen) supplemented with 5% donor horse serum (Sigma), 20 ng/ml epidermal growth factor (EGF; Sigma), 10 μg/ml insulin (Invitrogen), 100 μg/ml hydrocortisone (Sigma), 1 ng/ml cholera toxin (Sigma) and 100 U ml^−1^ penicillin and streptomycin (Invitrogen).

#### Human-derived samples

Patient-derived xenotransplanted (PDX) T-ALL cells and human-derived thymocytes were isolated as described previously (Akkapeddi *et al*, 2019; Silva *et al*, 2011). In all cases, informed consent was obtained in accordance with the Declaration of Helsinki and under institutional ethical review board approval. *Ficoll-Paque*^*™*^ *PLUS* (Thermo Fisher Scientific; 11768538) density gradient centrifugation was performed to separate dead cells, following standard methodology. Briefly, cells were resuspended in RPMI 1640 medium supplemented with 2mM L-glutamine, 1% PSA and 10% FBS. Diluted cells were gently added to a 15ml centrifugation tube containing 1/3 of the final volume until having two separate layers clearly defined of cells and Ficoll. The solution was centrifuged at 1000g 20 min at RT without the brake and viable cells were collected from the ring formed between the layer of Ficoll and medium. Leukemia cells were resuspended in medium, washed twice by centrifugation (800g for 7 min at RT) and counted.

### Immunofluorescence

Adherent cell lines were directly grown on glass coverslips. Suspension cell lines were pelleted and then re-suspended in 100 μL of 1× Dulbecco’s phosphate buffered saline (PBS) solution without Calcium and Magnesium (Biowest) and cytospinned onto slides using a Wescor Inc 7620 Cytopro^™^ Cytocentrifuge (500 rpm for 5 min at medium acceleration). Subsequently, both adherent and suspension cells were fixed using cold methanol for 10 min at −20 °C and incubated with blocking buffer (1× PBS containing 3% BSA and 0,1% Triton ×100) for 30 min at room temperature. Cells were then incubated for 2 hours at room temperature with the following primary antibodies: anti-Centrin-2 mouse clone 20H5 (1:1000, Milipore), anti-α-tubulin rat YL1/2 MCA776 (1:500, Serotec) and anti-CP110 rabbit (1:250, homemade (Jiang *et al*, 2012)) in blocking buffer. Cells were washed 3 times with 1X PBS solution and incubated for 1 hour at room temperature with the following secondary antibodies: anti-Ig G mouse conjugated with Alexa488 (Molecular probes), anti-Ig G rat conjugated with Alexa555 (Life technologies) and anti-Ig G rabbit conjugated with Alexa647 (Thermo Fisher), all diluted at 1/500 in blocking buffer. DAPI was also added to stain the DNA. Finally, the coverslips were washed 3 times with 1× PBS solution and mounted on slides using ProLong® Gold Antifade Reagent (Molecular Probes). The slides were kept 24 h at room temperature to allow the mounting media to cure. Mitotic spindles were analysed by using FIJI (ImageJ) software. Mitotic cells were identified by bright DAPI signal and by visualizing the spindle due to the α-tubulin staining. Only centrioles positive for Centrin-2 and CP110 were analysed and scored. At least 30 mitotic cells with centrosome amplification (cells with >4 centrioles) were quantified per cell line for the analysis of CA coping mechanism mechanisms. Figures were built using INKSCAPE software.

### *In vitro* drug testing

#### Chemicals

Recombinant human TGFβ1 (*240-B R&D Systems*) was stocked as 20 μg/ml in DMSO and AZ3146 (*SML-1427 Sigma Aldrich*) was stocked as 10 mM solution in 0.1% BSA, 4 mM HCl respectively. The appropriate amount of indicated solvents was employed for negative control conditions in each case.

#### TGFβ1 treatment

HS578T breast cancer cell line was thawed and cultured as mentioned in Table 5 in a 100 mm plate. The following day cells were transferred to 4 wells of a 6 well plate (25% confluency). Recombinant human TGFβ1 was added to two of the wells at a final concentration of 5ng/ml for 6 days to induce EMT. 72h after the seeding, cells were trypsinized and transferred to a p24 well plate with 13 mm glass coverslips placed at the bottom (6 replicates per condition). At the end of the 6 days of treatment, cells were fixed in ice cold methanol for 10 minutes at -20ºC. MCF10A-iPLK4 cells were thawed onto a 100mm plate. The following day cells were transferred to 4 wells of a 6 well plate (30% confluency). TGFβ1 was added to 2 of the wells at a final concentration of 5ng/ml. 48h later, Plk4 expression was induced with 2μg/ml in 1 control well and in 1 well treated with TGFβ1 and cells were incubated more 48h. Cells were then transferred to p24 well plates with 13 mm coverslips. Doxycycline induction was released and TGFβ treatment kept for more 48h, when cells were fixed for 10 minutes with ice cold Methanol at -20ºC (See flowchart in Fig. 2J). For all experiments, the supplemented medium together with TGFβ1 or the solvent was refreshed every other day. More than 50 and 100 cells with CA were scored per condition for HS578T and MCF10A-iPLK4 cell lines respectively and each experiment was repeated three times.

#### MPS1 Inhibitor (AZ3146) treatment

ALL cell lines (JURKAT, DND41, P12 and HBT-ALL) were seeded in a flat-bottom 96-well plate as 0.5×10^6^ cells/mL and HELA and RPE cell lines at seeding densities of 30% of confluency. 8 wells for condition were plated; either treated with 2.5 or 5 μM of AZ3146 inhibitor or DMSO. After 48 and 72 h, cell viability was assessed according to the manufacturer’s instructions (MTT ASSAY kit (Abcam ab211091). Absorbance was measured at 590 nm using the Multiskan Sky (ThermoFisher). Cell viability was also manually measured in ALL cell lines (JURKAT, DND41, P12 and HBT-ALL) by using the Handheld Automated Cell Counter Scepter ^™^ (Milipore). JURKAT, DND41, P12 (0.5×10^6^ cells/mL) and HPB-ALL (1×10^6^ cells/mL) were seeded in a flat-bottom 12-well plate. 5 μM of AZ3146 inhibitor or DMSO was added to the well and cell viability was assessed after 72h. The spindle phenotype was also analysed in all the ALL-cell lines. Each experiment was performed at least in triplicates and repeated twice.

### Image acquisition and analysis

Cell lines were observed on a commercial Nikon High Content Screening microscope, based on Nikon Ti. Images were acquired with an Andor Zyla 4.2 sCMOS camera, using a 100× 1.45NA oil immersion objective, DAPI + FITC + TRITC + Cy5 fluorescence filtersets and controlled with the Nikon Elements software. These images were deconvolved with the AutoQuantX3 software (MEDIACybernetics). All the images were taken as Z-stacks in a range of 7-9 μm, with a distance between planes of 0.2 μm.

### Immunoblotting

HS578T and MCF10A-iPLK4 cells were harvested and lysed with lysis buffer (50mM HEPES, 1mM EGTA, 1mM MgCl2, 100mM KCl, 10% Glycerol and 0.05% NP-40) with added protease inhibitors (Roche). Protein concentration was quantified using the Bio-Rad DC protein assay (25 μg loaded per well). Protein samples were resuspended in Laemmli buffer, separated on SDS-PAGE mini-protean precast gradient gels (BioRad REF) and transferred onto nitrocellulose membranes. Immunodetection was performed by incubation with antibodies against E-cadherin (1:1000; 3195S from Cell signaling), Vimentin (1:250; sc-373717), fibronectin (1:250; sc-18825) and N-cadherin (1:200; sc-59987) from Santa Cruz Biotechnology. GAPDH (1:1000; 2118S from Cell signaling) was used to normalize target protein abundance. After incubation with horseradish peroxidase–conjugated secondary antibodies (1:5000; Bethyl Laboratories), the immunolabeled proteins were detected by WesternBright^™^ ECL Luminol Reagent (Advansta) and developed by chemiluminescence (Amersham^™^ Imager 680, GE).

### NCI-60 metadata and molecular datasets

Metadata, including the information on doubling time, as well as gene and protein expression datasets for the NCI-60 cell lines were retrieved from CellMiner (https://discover.nci.nih.gov/cellminer/) (Shankavaram *et al*, 2009) and subset for the cell lines of interest. No data was available for the breast cancer cell line MDA-MB-468, which we therefore excluded from downstream analyses. Gene expression levels were retrieved in the form of average z-scores of intensities measured by five different microarray platforms (Reinhold *et al*, 2012). We removed the CNS cell line SF-539 from the dataset since it lacked expression data for approximately one third of the genes. Low variance genes were filtered out leaving 16,698 genes across 27 cell lines for analysis. We analyzed three different protein level datasets. 1) Log2-expression values for 52 cancer-associated proteins calculated by dose interpolation analysis from reverse-phase protein lysate arrays (Nishizuka *et al*, 2003). 2) Mass-spectrometry derived relative abundances (Gholami *et al*, 2013) that were filtered at the level of cell lines and individual proteins. First, the cell lines LC-HOP-62 and SK-MEL-28 were removed from the dataset as principal component analysis (PCA), performed with the *factoextra* R package, showed their global protein expression profiles to be extreme outliers. Then, proteins with data for less than 12 cell lines, i.e., half of the panel, as well as low variance proteins were removed, yielding a final dataset of 2,998 proteins across 26 cell lines for analysis. 3) Normalized protein levels determined by SWATH mass spectrometry were downloaded (Guo *et al*, 2019) and low variance proteins were filtered out, leaving 1,899 proteins across 28 cell lines for analysis.

### Development of a linear regression model for detection of centrosome clustering associated gene expression

To unravel which interesting aspects of the cancer cell lines correlate with clustering ability, we searched for genes whose expression levels correlate with CC capacity using the publicly available gene expression datasets of the NCI-60 panel.

For most genes, the Spearman correlation with CC was much more significant than expected by chance (determined by a Chi-squared goodness of fit test with correlation coefficients grouped into ten percentiles). Since the leukemia cell lines have a very distinct transcriptional profile (Fig. S1B) and a significantly stronger CC capability than cell lines from other cancer types (Fig. 1D), we reasoned that the correlation may partially reflect leukemia-specific gene expression. Indeed, repetition of the analysis without the leukemia cell lines brought the significance of most genes’ correlation with CC closer to what is expected at random. Similarly, the cell lines’ proliferation rates may be another confounding factor, since CC capability correlates with doubling time (Fig. S1C).

To test the likely confounding effect of the cell lines’ leukemia status and doubling time, individual linear regression was used to model the differential gene expression between leukemia and other cell lines, associated with doubling time or with CC-capability. The gene expression changes associated with the three effects were strongly correlated with each other and with the Spearman correlation coefficient from the previous analysis (Fig. S1D), indicating that the latter is not a suitable approach to identify genes whose expression is specifically associated with CC. To control for these confounding factors, we fitted a multivariate linear model to the expression of each gene including, as independent variables, CC capability, leukemia status and doubling time. Albeit not fully decoupling them, this approach dramatically reduces the correlation between the different effects (Fig. S1E-F) and provided a more accurate quantification of the independent association between CC and each genes’ expression levels across all cell lines. We tested for multicollinearity of the variables in the linear model by calculating the variance inflation factor (VIF) using the *vif* function within the *car* R package. The calculated VIF was for 1.9 for CC, 1.8 for the leukemia status and 1.4 for the doubling time, indicating that collinearity between the independent variables is small enough to separate their effects in the model. PCA was performed with the *factoextra* R package. Spearman correlation was calculated using the *cor*.*test* function and the Chi-squared goodness of fit test was done using the *chisq*.*test* function from the *stats* R package. Linear regression was performed with the *limma* R package (Ritchie *et al*, 2015). Correction for multiple testing was performed by calculating the false discovery rate (FDR) with the *p. adjust* function from the *stats* R package.

### Gene Set Enrichment Analysis

The curated lists of gene sets “Hallmark gene sets” (n = 50) and “gene ontology (GO) cellular component” (n = 580) were retrieved from the Broad Institute’s Molecular Signatures Database v6.2 (Subramanian *et al*, 2005; Liberzon *et al*, 2015) (Table 2). These gene sets are representative of major cellular components and pathways. Moreover, a list of gene sets representative of signalling pathways was retrieved from SPEED (Parikh *et al*, 2010).

Furthermore, we generated curated lists of centriole duplication factors, CA genes (Ogden *et al*, 2017) and a list of known CC-factors generated from literature research (Drosopoulos *et al*, 2014; Kwon *et al*, 2008; Leber *et al*, 2010), as well as several subsets of this list with genes grouped by their function (Table 03). Gene sets for epithelial and mesenchymal features were also retrieved from the literature and manually curated ((Table 03) (Lee *et al*, 2006, 2006; Zhao *et al*, 2015; Mak *et al*, 2016; Rokavec *et al*, 2017; Gröger *et al*, 2012; Lee & Nelson, 2012)).

Gene Set Enrichment Analysis (GSEA) (Subramanian *et al*, 2005) was used to identify sets of genes from those lists over-represented (“enriched”) amongst those more associated with CC. In each analysis, genes were ranked by their t-statistic of association with CC. This analysis was carried out in R using the *piano* package (Väremo *et al*, 2013), including 10,000 gene list permutations to estimate significance and FDR correction for multiple testing, was done with all gene sets at once. The code for running enrichment score plots was adapted from https://github.com/ctlab/fgsea/blob/master/R/plot.R.

### Gene expression analysis during breast cancer malignant progression

We compiled published transcription profiles of 893 samples from different progression stages of breast cancer: normal breast epithelium (N), simple hyperplasia (SH), atypical hyperplasia (AH), ductal carcinoma in situ (DCIS) and invasive ductal carcinoma (IDC) (Table 04). After data pre-processing using *fRMA* (McCall *et al*, 2010), all datasets were merged and expected batch effects were removed using *ComBat* (Johnson *et al*, 2007). PCA was performed with the *factoextra* R package (Lê *et al*, 2008) (Fig S3A-C). We performed differential gene expression analyses between samples of each cancer progression stage and normal breast tissue samples and tested the association with centriole duplication (manually curated list) and centrosome clustering (compiled from 3 RNAi-screens: (Drosopoulos *et al*, 2014; Kwon *et al*, 2008; Leber *et al*, 2010) genes through GSEA, as described above.

### Survival analyses

Survival analyses were performed using gene expression and clinical data from multiple breast cancer datasets with KM-plotter (http://kmplot.com/analysis/index.php?p=service&cancer=breast, (Győrffy, 2021)). We tested the association between overall survival and the combined expression of genes involved in centriole duplication and function (selected genes: *PLK4, BRCA1* and *AURKA*, (Qi *et al*, 2022; Magnaghi-Jaulin *et al*, 2019; Tkach *et al*, 2022; Moyer & Holland, 2019)) and regulators of CC (*STK10, DBR1*, and *KIFC1*, (Pannu *et al*, 2015b; Vitre *et al*, 2020; Chavali *et al*, 2016; Kwon *et al*, 2008)) both independently and combined. Patients were split by the median combined expression of the gene set (combined by averaging the expression of the selected genes) and significance was assessed with the log rank test.

### Other statistical analyses

Other statistical analyses were conducted using GraphPad Prism (Version 5.01. 2007 GraphPad Software, Inc.). A two-tailed 5% significance level was considered as statistically significant (*p* < 0.05). Statistical comparisons were assessed by unpaired two-tailed *t*-test and ordinary two-way ANOVA (Tukey’s multiple comparison test).

## ACKNOWLEDGEMENTS

We would like to acknowledge all members of the M.B-D. lab for critical reading and fruitful discussions on the manuscript. We are thankful to Manuel Théry for helpful discussions, David Pellman for discussions and motivating us for this study, and all the people that provided us with the cell lines. We thank Paulo Duarte for technical support. The authors would like to acknowledge the IGC Light Microscopy facility members, especially Mária Hanulová, José Marques, and Gabriel Martins for equipment availability and helpful support. This work was supported by FCT (Fundação para a Ciência e Tecnologia) – grants HMSP-CT/SAU-ICT/0075/2009 and PTDC/BIM-ONC/6858/2014.

## AUTHOR CONTRIBUTIONS

N. Moreno-Marín and M. Bettencourt-Dias investigated, did the formal analysis and curated the data, conceptualized and wrote the original draft; G. Marteil, with the help of K. Dores, designed and performed the screening on the CA-coping mechanisms. N. C. Fresmann and B.P. de Almeida, with the supervision of N. Morais, ran the bioinformatic analysis and provided the plots associated. J.T. Barata and R. Fragoso provided the ALL PdXs and critical ideas regarding leukemia. J.Vas and J. Pereira-Leal ran the initial analysis related to breast cancer. S. Godinho helps to conceive some of the experiments. All the authors reviewed and discussed the manuscript.

---

**Fig. S1.**
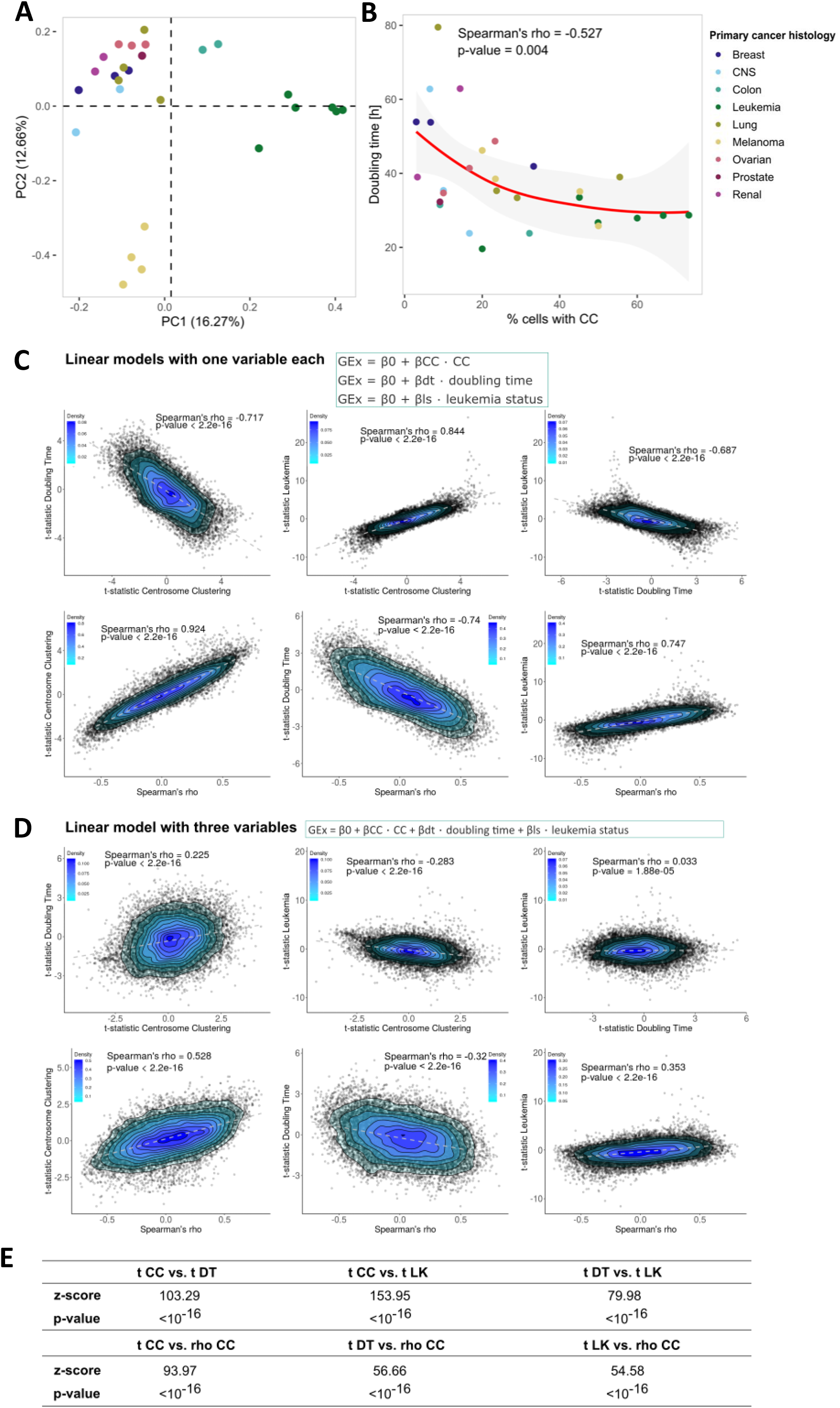
A linear model partially decouples the effects of centrosome clustering, tissue of origin and doubling time on gene expression. **(A)** Cell lines derived from leukemia show a transcriptional profile that is very distinct from the rest of the tissues represented in the NCI60 panel, as shown by PCA analysis of publicly available transcriptomic data (see Materials and Methods). The 27 cell lines with sufficient available gene expression data are shown and plotted by the first two components of a PCA using the transcriptomic data. Cell lines are colored by tissue of origin shown in B. **(B)** The percentage of cells that cluster centrosomes (X-axis) is inversely correlated with the publicly available cell doubling time of 28 of the cell lines (Y-axis, Spearman correlation with p-value < 0.05). The cell line MDA-MB-468 was excluded due to a lack of information on the doubling time. The grey shade around the red *Loess line* represents the 95 % confidence interval. Color legend as in A. **(C)** Centrosome clustering, tissue of origin (leukemia vs others) and the doubling time of the cells are strongly correlated. Upper panels: Gene expression (GEx) is linearly modelled by one variable β_*effect*_(centrosome clustering (CC), tissue of origin (leukemia or others (ls)) or doubling time(dt)), apart from baseline β_0_ (model’s intercept), as per the equations above. The t-statistics of differential expression associated with the corresponding effects are plotted against each other (left: centrosome clustering (CC) vs. doubling time (dt), middle panels: centrosome clustering vs. leukemia (tissue of origin), right: dt vs. leukemia). Lower panels: t-statistics of differential expression for each of the effects plotted against the Spearman’s correlation coefficient (rho) between centrosome clustering and gene expression. **(D)** Gene expression was linearly modelled by the three independent variables together. The t-statistics of differential expression associated with the individual effects were plotted against each other and are less correlated than those for the individual models shown in (C). Albeit not fully decoupling them, this approach dramatically reduces the correlation between the different effects, providing a more accurate quantification of the independent association between CC and each genes’ expression levels across all cell lines. **(E)** The differences between homologous correlations shown in (C) and in (D) were calculated in z-values, using the Fisher r-to-z transformation. All pairs of correlations are significantly different (p < 10^−6^).

**Fig. S2.**
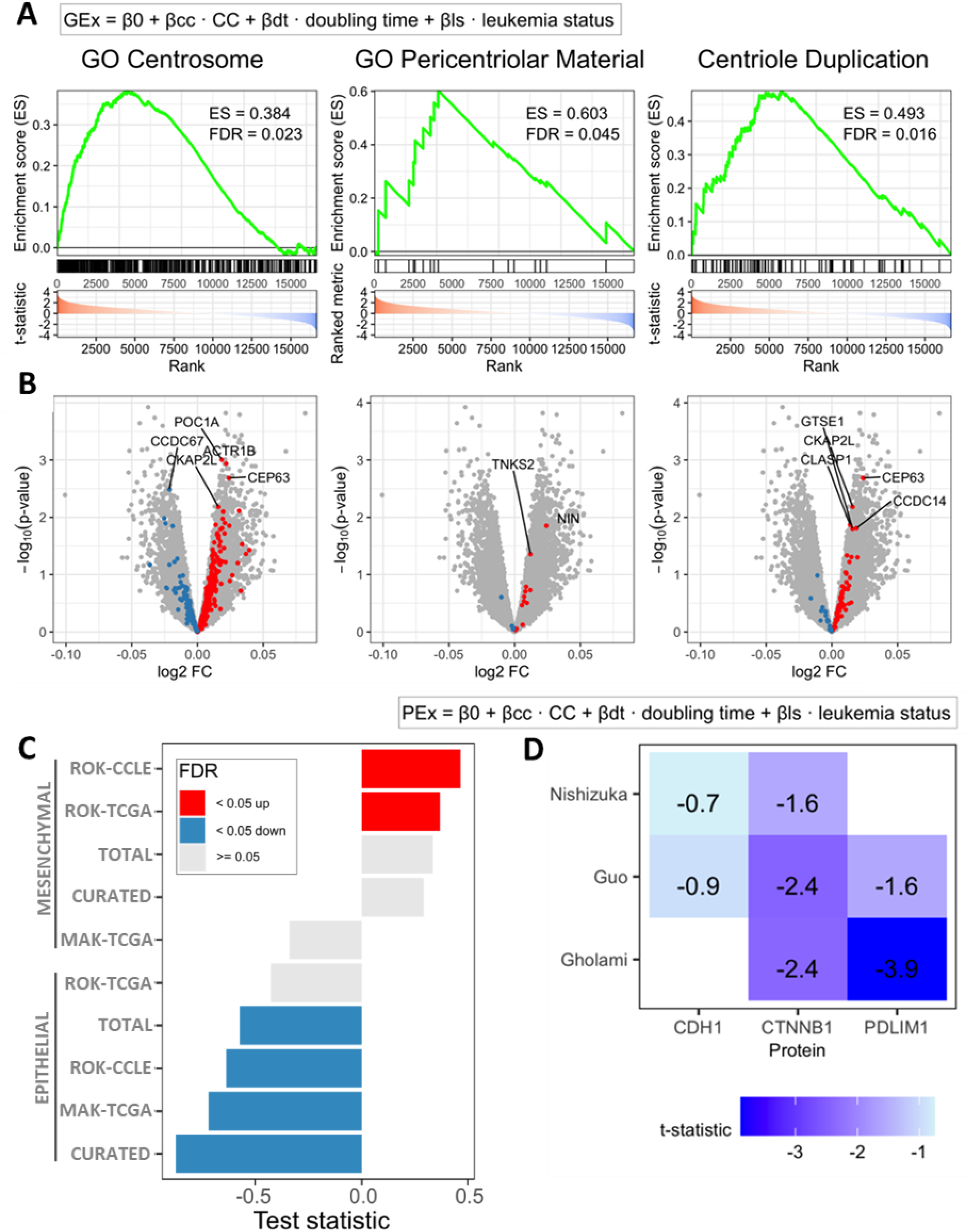
Centrosome clustering is associated with the loss of epithelial properties and the EMT in the NCI60 panel. **(A)** GSEA of centriole-associated gene sets with genes ranked by their association with CC in the linear model. GSEA peak enrichment scores (ES) and false discovery rates (FDR) are shown. Genes are ranked by their t-statistic of association with CC and plotted on the X-axis. Genes that belong to the gene set of interest are indicated by vertical black bars. The enrichment score is plotted in green. **(B)** Volcano plots of differential gene expression associated with CC. Genes that belong to the gene sets in (A) are highlighted according to whether they are negatively (blue) or positively (red) associated with CC. **(C)** Barplot showing GSEA enrichment scores for all gene sets comprising mesenchymal or epithelial genes. The FDR was calculated using the full set of gene sets used in this study. Gene sets with significant genes over or under-expressed in association with CC (FDR < 0.05) are represented in red and blue, respectively. **(D)** T-statistic for association of protein expression with CC, linearly modelled according to the equation at the top using three independent datasets (Gholami *et al*, 2013; Nishizuka *et al*, 2003; Guo *et al*, 2019) shows consistent, albeit not significant, negative association of E-cadherin (CDH1), β-catenin (CTNNB1) and PDLIM1 levels with CC (Table 1). PDLIM1 downregulation has been associated with metastatic potential in colorectal cancer through destabilization of the E-cadherin/β-catenin complex (Chen *et al*, 2016)

**Fig. S3.**
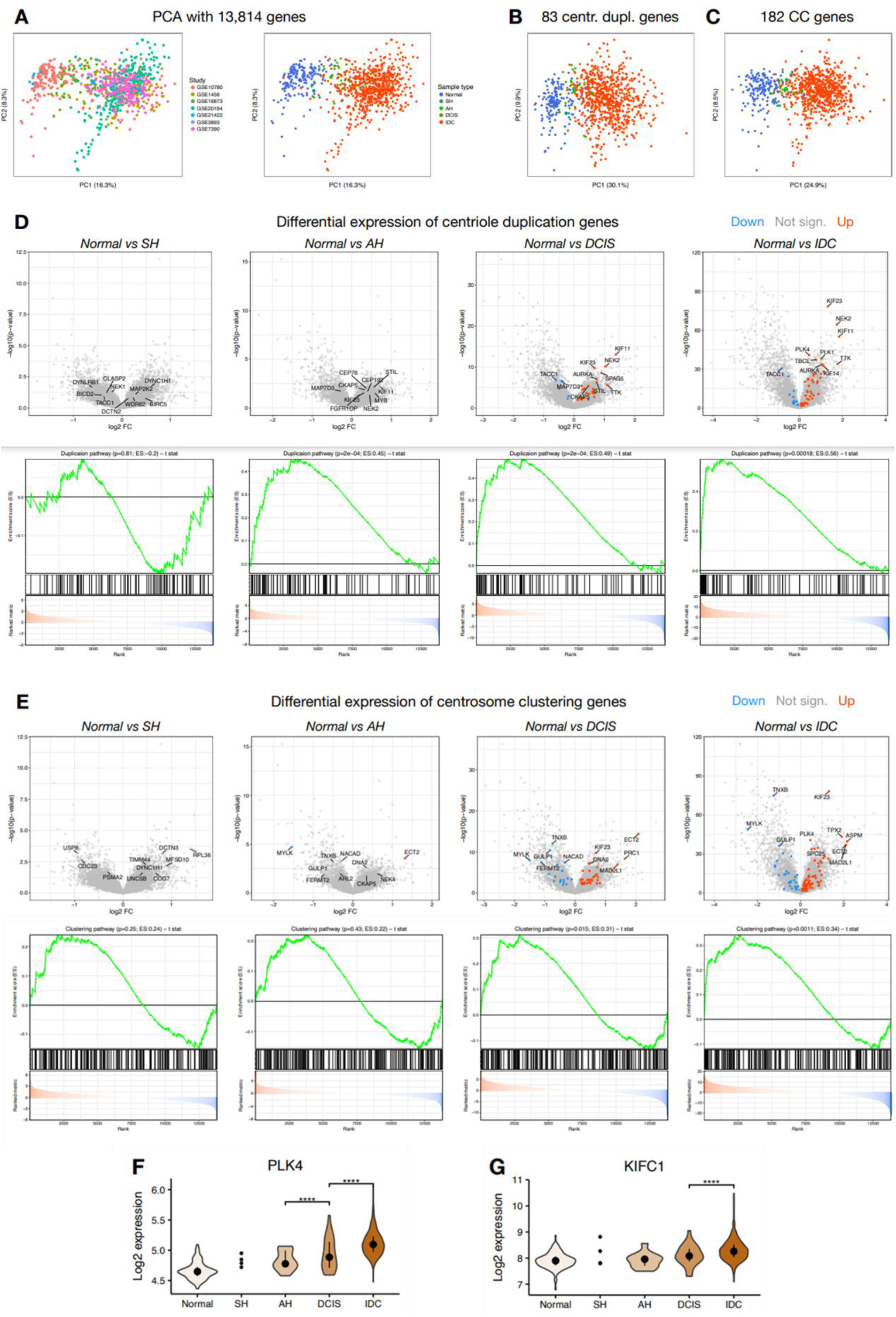
Centrosome clustering is associated with breast cancer malignant progression. **(A-C)** Principal Component Analysis using **(A)** all 13,814 genes, **(B)** 83 centriole duplication genes and **(C)** 182 centrosome clustering genes. Principal Components (PC) 1 and 2 are shown. Samples are colored by study of origin or sample type. **(D-E)** Differential expression of **(D)** centriole duplication genes and **(E)** centrosome clustering genes between normal samples and each type of tumor samples. Top: volcano plots of differential expression with genes over or under-expressed compared to normal samples (FDR < 0.05) represented in red and blue, respectively. Top 10 genes are labeled. Bottom: Gene Set Enrichment Analysis (GSEA) of genes ranked by the differential expression t-statistics. GSEA p-values are shown. SH, simple hyperplasia, AH, atypical hyperplasia; DCIS, ductal carcinoma *in-situ*; IDC, invasive ductal carcinoma. **(F-G)** Violin plots of log2 expression of **(F)** PLK4 and **(G)** KIFC1 genes across samples from different breast tumor types. ^****^p-value < 0.0001 (Wilcoxon rank-sum test). Black points and lines represent the median +/-upper/lower quartiles. N, normal; SH, simple hyperplasia, AH, atypical hyperplasia; DCIS, ductal carcinoma *in-situ*; IDC, invasive ductal carcinoma. For SH, values for individual values are shown because violin plots require a minimum sample size of 6.

